# A Single Chromosome Strain of *S. cerevisiae* Exhibits Diminished Ethanol Metabolism and Tolerance

**DOI:** 10.1101/2020.08.22.256727

**Authors:** Tyler W. Doughty, Rosemary Yu, Lucy Fang-I Chao, Zhongjun Qin, Verena Siewers, Jens Nielsen

## Abstract

This study characterized the growth, metabolism, and transcriptional profile of a *S. cerevisiae* strain with a single large chromosome that was constructed via successive chromosomal fusions. The single chromosome strain exhibited a longer lag phase, increased doubling time, and lower final biomass concentration compared with a wildtype strain when grown on YPD. These phenotypes were amplified when ethanol was added to the medium or used as the sole carbon source. RNAseq analysis showed diminished induction of genes involved in diauxic shift, ethanol metabolism, fatty-acid ß-oxidation, and methylglyoxal catabolism during growth on ethanol compared to the reference strain. Enzyme-constrained metabolic modeling predicted that decreased flux through these poorly induced enzymes results in diminished ATP formation and decreased biomass accumulation observed. Together, these observations suggest that switch-like control of carbon source dependent gene expression in *S. cerevisiae* requires genome arrangement into multiple chromosomes.

## Introduction

The nuclear genetic code of eukaryotic organisms is arranged as linear chromosomes that facilitate organization and protection of DNA^1–4^. Although chromosomes were observed as early as 1842, the characterization of the sequence and function of centromeres, telomeres, and autonomous replication sequences occurred much later^4,5^, as technology like DNA sequencing became more readily available^6^. Additional progress arose from disruption of chromosomal substructures, followed by characterization of mutants, and forward engineering. Examples of this paradigm include the sequencing, characterization, and forward engineering of mitotically segregated plasmids^2^, as well as the design of artificial chromosomes in yeast (YACs)^7,8^. Recent advances in high throughput sequencing have enabled sequencing and assembly of whole genomes, analysis of transcriptomes, and reconstruction of 3D chromosome structure^9–11^. The emergence of these tools coincides with advances in CRISPR-Cas9 genome editing^12^, which has enabled scientists to disrupt chromosomal organization by creating novel genomic structures. Analysis of these novel mutant organisms may shed light on previously uncharacterized fundamental questions by creating novel genomic structures.

Two recent reports applied the aforementioned technologies to create and analyze *S. cerevisiae* strains with one^13^ or two^14^ chromosomes via successive breaks of chromosome ends followed by repair/fusion^13,14^. These end-to-end fusions are similar to events that can occur naturally during vast evolutionary timescales^15,16^, and enabled novel analyses of meiosis^14^ and chromosomal folding^13^ in the context of a few large chromosomes versus sixteen smaller chromosomes. Intriguingly, the single chromosome strain from Shao et al. 2018 exhibited similar glucose phase growth rates and gene expression compared to the wildtype, despite significant changes in 3D chromosomal organization and interchromosomal interactions^13^. Additional comparative studies of reference and chromosome fusion strains to determine their phenotypic differences may shed light on the causes and consequences of chromosome organization, which will inform evolutionary biology and enable the design of synthetic chromosomes.

In this work, we investigated growth, gene expression, and metabolism of the single chromosome yeast strain from Shao et al. 2018^13^ compared with a reference strain during glucose and ethanol phase growth. We observed decreased biomass accumulation, decreased viability in the ethanol phase, and a dose dependent sensitivity to ethanol compared with the reference strain. Transcriptomics and metabolic modeling suggest that these phenotypes were influenced by improper activation of non-fermentable carbon source utilization and diauxic shift genes. We hypothesize that the gene expression regime that enables growth and survival on non-fermentable carbon sources is dependent on chromosomal organization, which is disrupted in the single chromosome strain.

## Results

### Profiling Single Chromosome Strain Growth

To characterize perturbations in growth and/or metabolism, we performed batch fermentations with triplicate bioreactors for the reference strain (*S. cerevisiae* strain BY4742) or the chromosomal fusion strain SY14, which has a single large chromosome instead of sixteen distinct chromosomes (Figure 1). Analysis of the CO_2_ evolution rate of the gas emitted from fermenters suggested that SY14 had an increased lag time prior to exponential growth on glucose (Figure 1A). In addition, the doubling time during growth on glucose was increased by 8% for cultures of SY14 (120minutes) compared with BY4742 (111minutes) (Figure 1C). Biomass accumulation, monitored via OD_600_, was diminished and became more apparent in the later stages of growth, culminating in 28% less biomass after 48hours of growth (Figures 1B and 1D). Shake flask experiments showed a similar decrease in final biomass after 10 days (Supplemental Figure 1). Despite a longer lag phase, SY14 cultures exhibited similar profiles of carbon source uptake, including complete uptake of glucose and production, followed by consumption of ethanol (Figure 1E-I). Together, these findings indicate that SY14 exhibited a delay in growth after inoculation, had decreased glucose phase growth, and accumulated less biomass than the reference strain.

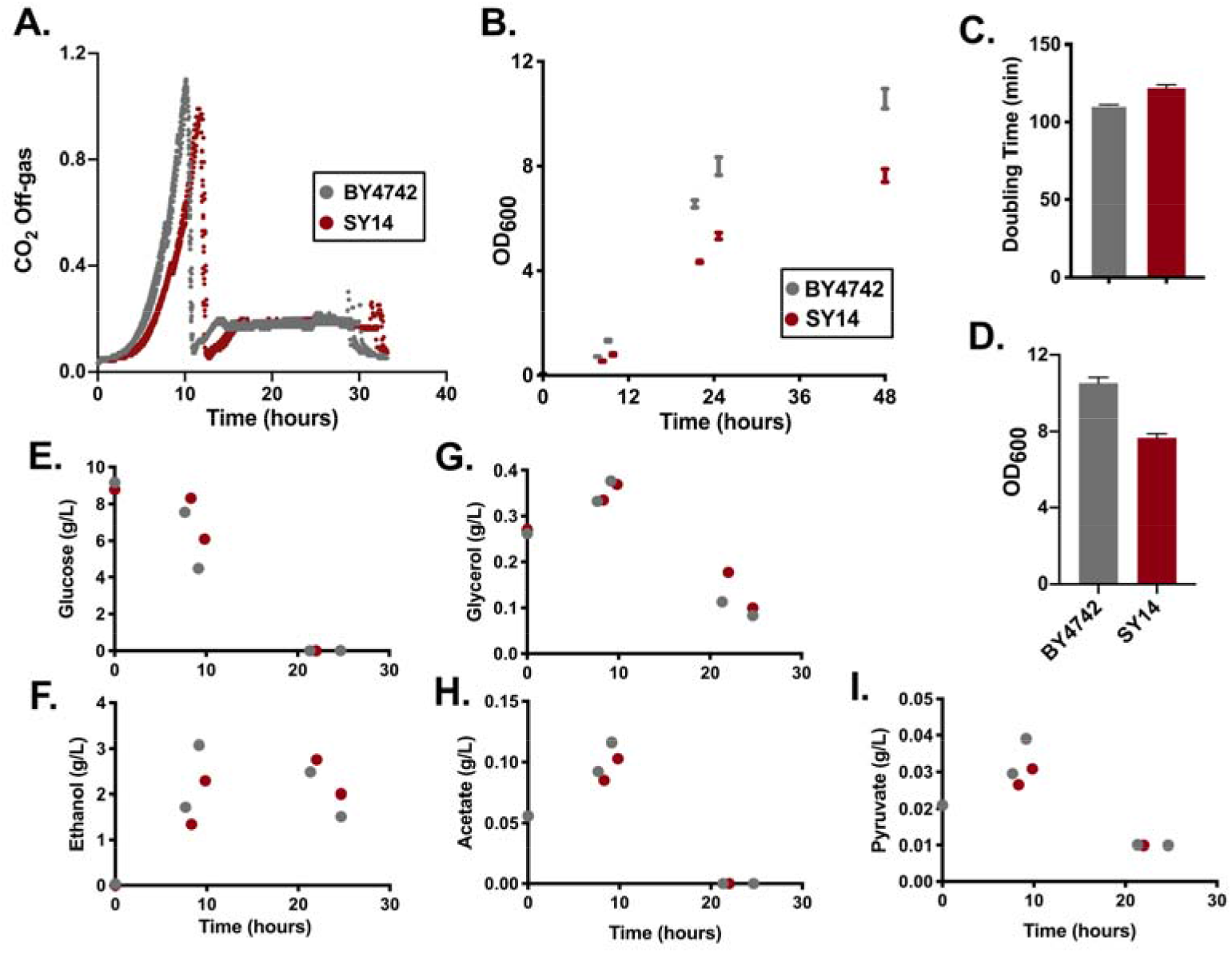
Batch Cultivation of a Single Chromosome Yeast Strain. Reference (BY4742) and single chromosome (SY14) strains were analyzed via batch fermentation to monitor CO_2_ evolution **(A)** and biomass accumulation **(B)**. Maximum glucose-specific doubling time **(C)** and the final OD_600_ at 48hours **(D)** were measured. **E-I**. HPLC was used to monitor media composition at various timepoints.

### The Single Chromosome Strain (SY14) Exhibits Impaired Growth on Non-Fermentable Carbon Sources and is Sensitive to Ethanol

The results presented in Figure 1 warranted further analysis of the various stages of growth for SY14 and the reference strain. Similar to the fermentation results, microplate growth assays showed that cultures of SY14 exhibited a longer lag phase, increased doubling time during growth on glucose, and lower final biomass yield than BY4742 (Figure 2A). Notably, the magnitude of these differences was larger for lag phase and final biomass than for glucose doubling time in microplate assays and fermenters. These differences suggested that SY14 cultures might struggle to grow on, and emerge from growth on non-fermentable carbon sources. To test this, we plated strains on glucose (YPD), ethanol (YPE), and glycerol (YPGly) plates (Figure 2B). The results showed that growth of SY14 was diminished compared to BY4742 on non-fermentable carbon sources, but was similar on glucose. These phenotypes did not appear to be due to oxidative stress that might occur during growth on non-fermentable carbon sources, as addition of 3mM H_2_O_2_ did not disproportionately influence SY14 doubling time or lag phase duration (Supplemental Figure 2). Further, total protein levels (Supplemental Figure 3A), as well as ribosomal RNA expression and processing were similar in wildtype and SY14 strains (Supplemental Figure 3B).

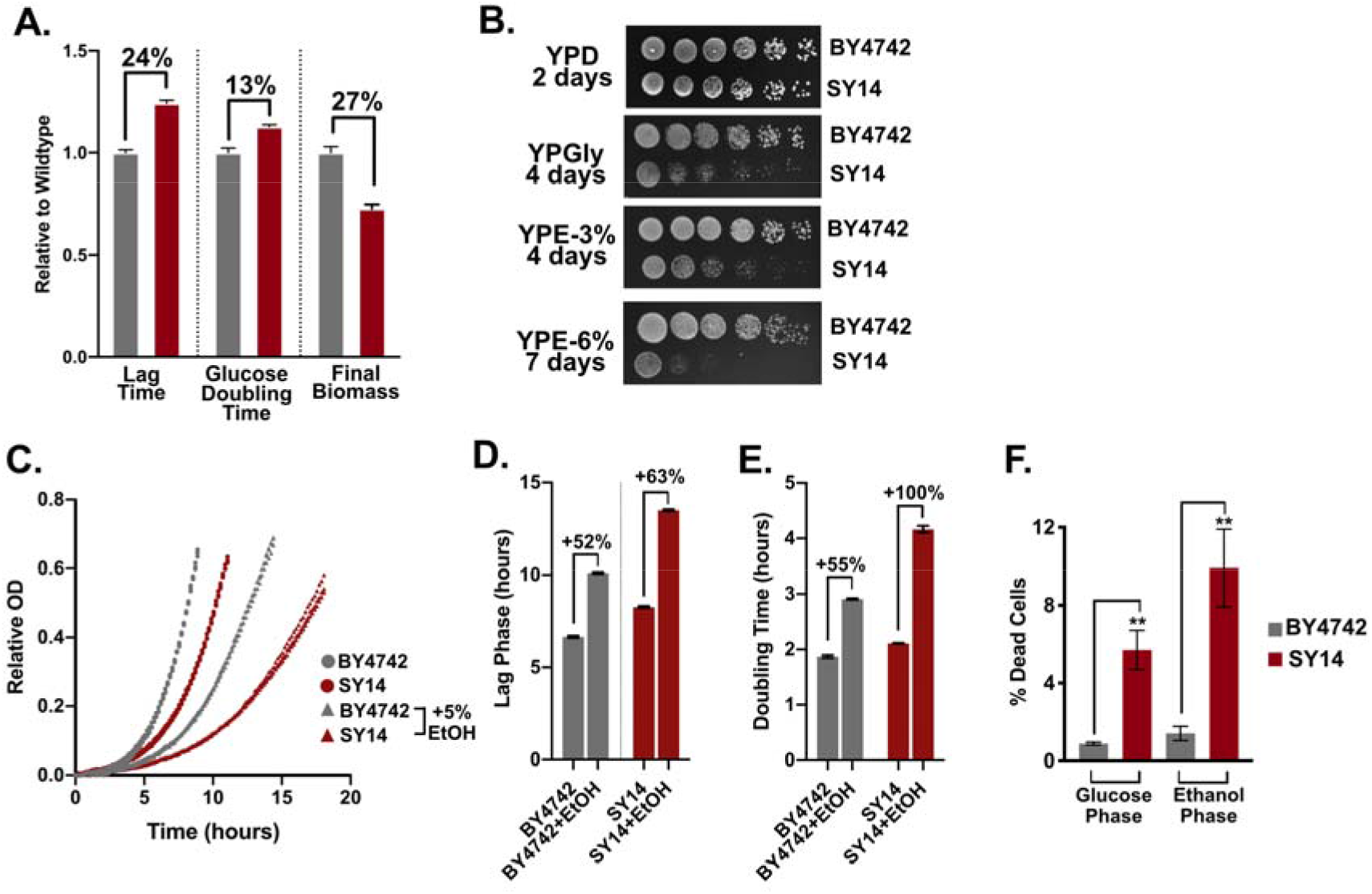
The Single Chromosome Yeast Grows Slowly on Non-fermentable Carbon Sources and is Sensitive to Ethanol. **A**. Single chromosome and reference strain growth was monitored to determine the time to reach an OD_600_ of 0.25 (lag phase), maximum glucose growth, and final biomass at 48 h. **B**. Growth on YP plates with varying carbon sources. Glucose phase growth in YPD +/-5% ethanol growth curves **(C)** and doubling time **(D). E**. Reference (BY4742) and single chromosome (SY14) strains were analyzed for cell death using propidium iodide staining.

**Figure 3.**
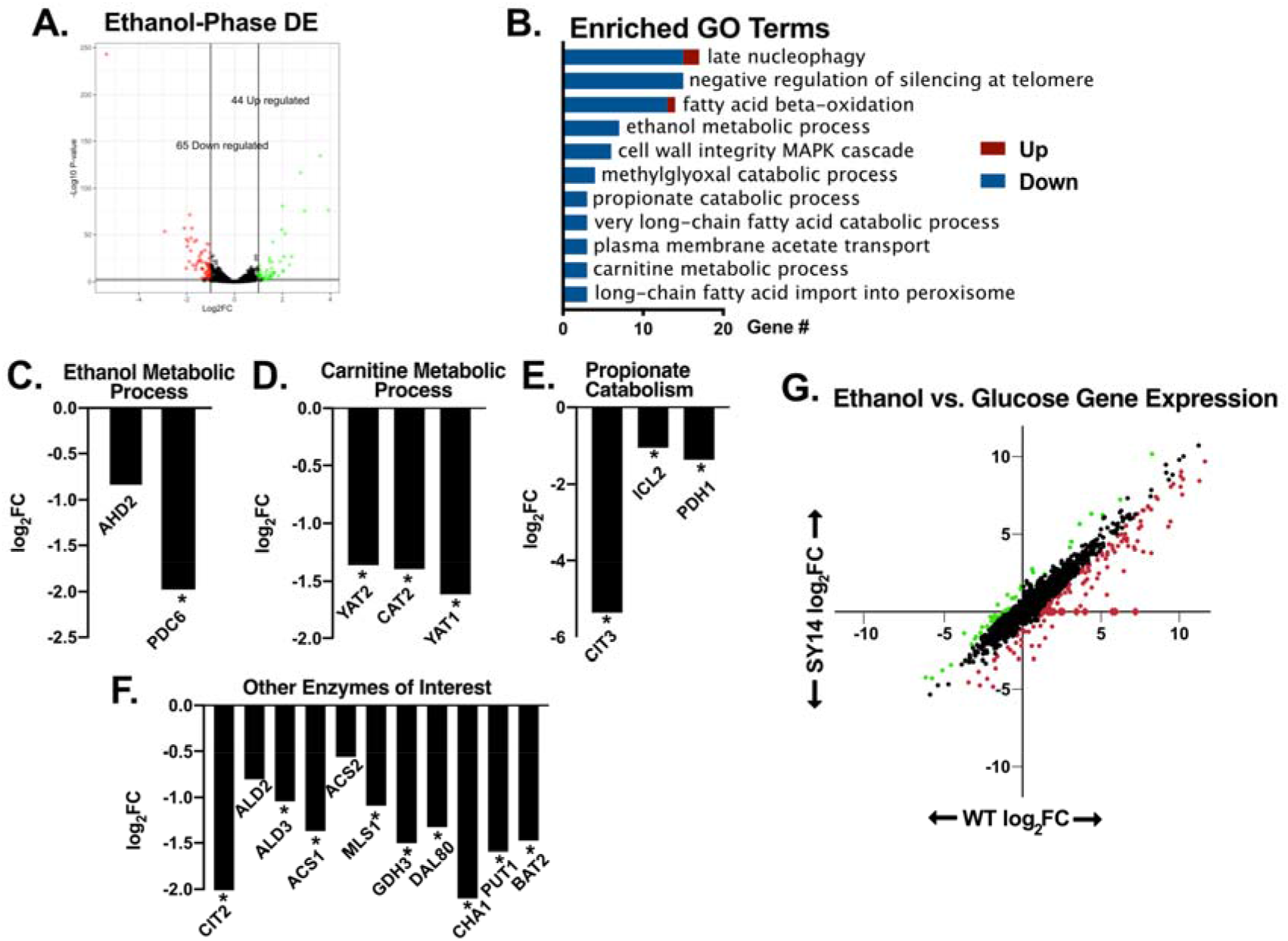
Ethanol-Phase RNA-seq Shows Decreased Induction of Diauxic Shift Genes in the Single Chromosome Strain. **A**. A volcano plot showing the differentially expressed genes in SY14 versus control after 20 h of growth. **B**. Downregulated genes from ethanol-phase RNAseq were assessed for enriched GO terms for strain SY14. **C-F**. Individual differential expression results are shown for select GO terms and genes. * indicates log_2_FC <-1 FDR<0.01. **G**. Gene expression changes between glucose and ethanol phase cultures of WT and SY14 are shown for comparison. Genes that exhibited increased (green) or decreased (red circles) in SY14 compared to wildtype are highlighted.

The results in Figure 2B showed that growth for SY14 was particularly diminished in the presence of 6% ethanol. To test for ethanol sensitivity, we cultured SY14 and BY4742 in YPD (glucose) media +/-5% ethanol (Figure 2C). The lag phase after inoculation was longer for SY14 with 5% ethanol (Figure 2D), and the doubling time during growth on glucose increased by 55% for BY4742 and 100% for SY14 (Figure 2E). These findings suggest that SY14 is sensitive to ethanol, even in the presence of glucose. This sensitivity may influence the observed increase in cell death in the SY14 background (Figure 2F).

### SY14 Exhibits Decreased Expression of Diauxic Shift Related Genes in the Ethanol Phase

Analysis of transcriptomic measurements during growth on glucose discovered relatively few differentially expressed genes in the SY14 background (53 genes) (Supplemental Figure 4A, Supplemental Table I). The number of differentially expressed genes is intriguing as chromosomal fusion drastically altered genome arrangement and disrupted many interchromosomal interactions, which are important for gene regulation in higher eukaryotes^17–19^. Furthermore, chromosomal fusion removed the majority of telomeres and centromeres which have previously been shown to influence gene silencing^13,20^. However, these gene expression results may not encapsulate the deficiencies of the strain during growth on non-fermentable carbon sources or in the presence of ethanol (Figure 2). To further understand these phenotypes, we performed RNA-seq to compare gene expression between the reference (BY4742) and single chromosome (SY14) strains during growth on ethanol following a glucose batch phase and the diauxic shift (Figure 3A). This analysis resulted in identification of a modest number of differentially expressed genes (109). Interestingly, genes with significantly lower expression in SY14 were enriched for functions related to growth on non-fermentable carbon sources (Figure 3B). Specifically, SY14 exhibited lower gene expression for enzymes involved in ethanol, carnitine, propionate, and fatty acid metabolism (Figure 3C-F), all of which enable *S. cerevisiae* to generate ATP after glucose depletion. The diminished gene expression observed might predict diminished growth post diauxic shift, which was observed in Figure 2B.

**Figure 4.**
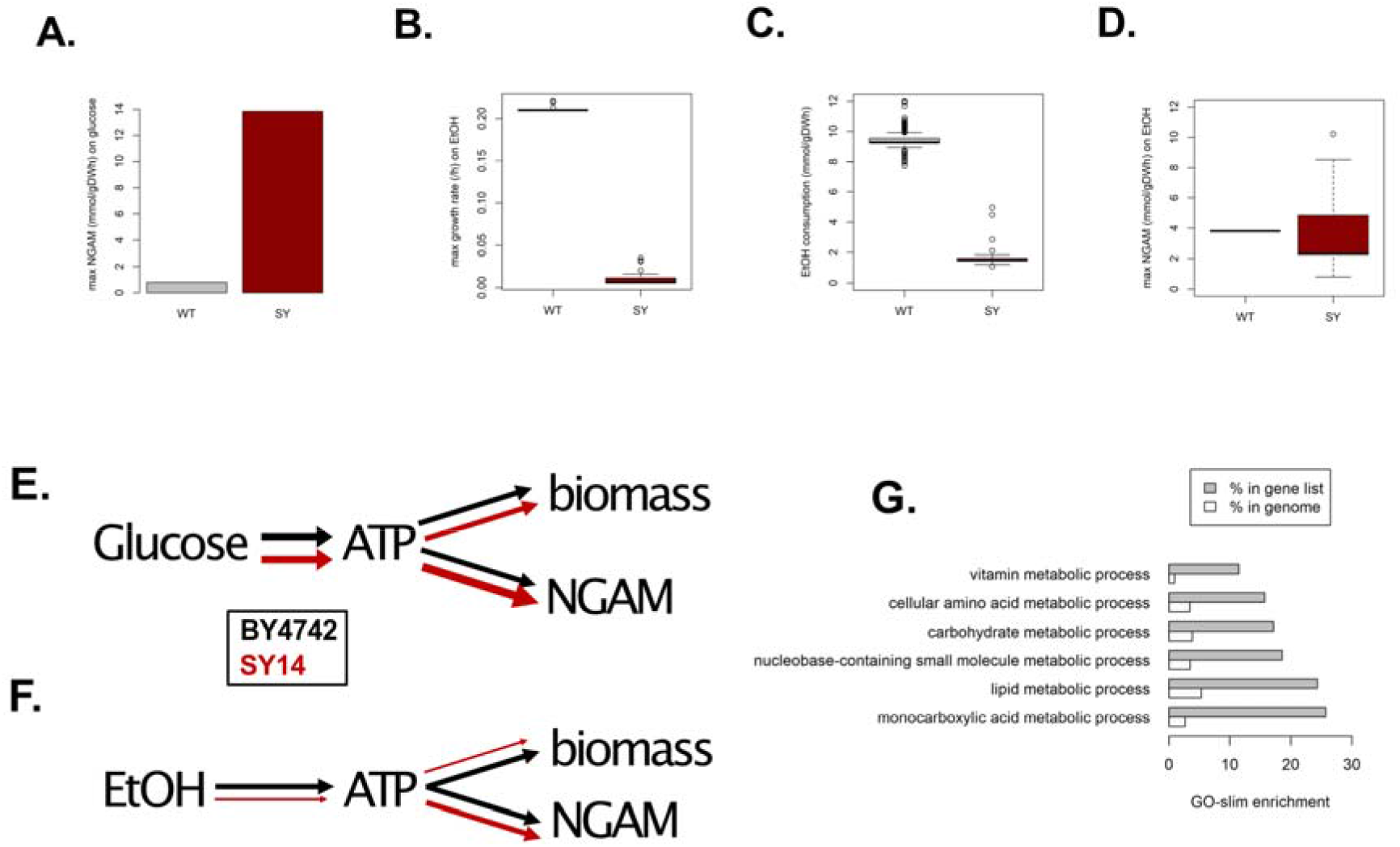
Enzyme-constrained genome scale modeling (ecGEM) predicts high NGAM on glucose and poor ATP generation on ethanol for the Single Chromosome Strain. A. An ecGEM was constructed to predict wildtype and SY14 ATP expenditure on non-growth associated maintenance (NGAM) when using glucose as the sole carbon source. A model reflecting the disrupted ORFs and differentially expressed genes was constructed to model growth on EtOH as the sole carbon source (B), ethanol consumption rates in SY14 (C), and max NGAM (D). Diagrams representing the growth defects in SY14 when using glucose (F) or EtOH (G) as the sole carbon source. Line thickness is indicative of relative flux for reference (black) and SY14 (red). G. Enzymes predicted to rescue growth defect of SY14 on ethanol were enriched for GO terms related to energy generation.

The diminished activation of the aforementioned genes was not associated with changes in sequence of the ORF, promoters, or terminators, with one exception, *CIT3*, which encodes a mitochondrial citrate and methylcitrate synthase. The promoter and 5’ coding region of CIT3, a gene known to be involved in propionate metabolism^21^, was removed during construction of the SY14 strain. Reintroduction of CIT3 via a plasmid did not alter the growth or ethanol tolerance of SY14 (Supplemental Figure 5), which might be expected as this gene was shown to encode a minor isoform of citrate synthase^21^. Further analysis of the RNAseq data identified several subtelomeric genes that were not expressed in the single chromosome strain (e.g. HSP33, PAU4, and AAD4), analysis of the SY14 genome sequence showed that these genes were likely removed during chromosomal fusion (Supplemental Figure 6). The majority of these deleted genes lacked a functional description (54%), or were members of the duplicated gene families PAU, COS, and AAD (24%). These genes were not amongst gene sets known to be essential^22^, associated with slow growth^22^, or known transcription factors^23^. Further, as a group, these deleted genes represented a small percentage of the total RNAseq reads in the wildtype strain during glucose (0.10%) or ethanol (0.16%) phase.

Next, we compared gene expression between glucose phase and ethanol phase for each strain to understand the transition between growth on different carbon sources. This analysis showed that several genes that were proximal to the remaining telomeres were upregulated during ethanol phase in the reference strain, but were not upregulated in the single chromosome strain (Supplemental Figure 7), suggesting that SY14 has increased subtelomeric silencing. Further analysis showed that several metabolic genes that were not telomere proximal exhibited lower induction during ethanol phase in SY14 compared to wildtype strains. We refer to these genes as poorly induced, as they are significantly upregulated (log_2_FC>1 FDR<0.01) in the wildtype strain upon transition from glucose to ethanol phase, but were at least two-fold less induced in the SY14 background compared to the reference strain (Figure 3G red dots). The 111 genes that were poorly induced in SY14 represented 3.56% of all RNAseq reads in ethanol phase samples, in contrast, these genes accounted for 7.1% of reads amongst reference samples. These data suggest that the significantly decreased ethanol phase expression of metabolic genes like *ACS1, YAT1, YAT2, CIT2, PDC6*, and *ADH2* in Figure 3C-F was due to a failure to upregulate these genes after the transition from glucose to non-fermentable carbon source growth in the SY14 background. The similarity of the functional annotations of the poorly induced genes in SY14 may indicate disruption of a global mechanism for regulating non-fermentable carbon source gene expression.

### Metabolic Modeling of SY14 Predicts an ATP Bottleneck During Ethanol Growth

The data in Figure 3 suggest that 248 enzymes were not properly upregulated in the SY14 background during growth on ethanol, which represents 26% of all metabolic enzymes in *S. cerevisiae*^24^. These genes and/or the subtelomeric genes that were disrupted during chromosomal fusion (Supplemental Figure 6), might explain the growth phenotype of the SY14 strain. To further understand how differentially expressed or disrupted genes might influence glucose and non-fermentable carbon source growth, we constructed enzyme-constrained Genome-scale Metabolic Models (ecGEM) for both reference and SY14 strains^25^, using downregulated and deleted genes to constrain enzyme usage in SY14 relative to the reference. As SY14 and reference cells exhibited remarkably similar metabolic profiles (Figure 1) and gene expression profiles (Supplemental Figure 4) in glucose-phase growth, the minor growth defect of SY14 pointed to an increased ATP expenditure for non-growth associated maintenance (NGAM), indicating that resources were being diverted to deal with stress (Figure 4A).

In contrast, modeling growth on ethanol as the carbon source showed that differentially expressed metabolic enzymes drastically limited the ability of SY14 to grow (Figure 4B) and utilize ethanol (Figure 4C). Of note, the calculated maximum ATP expenditure on NGAM was comparable between wildtype and SY14 during growth on ethanol (Figure 4D), indicating that a bottleneck in ATP generation from ethanol underlies the reduction in biomass formation for SY14. Together, this analysis suggested that when using glucose as a carbon source, the growth defect in SY14 cells arose from an increased ATP expenditure to handle stress (Figure 4E). Conversely, the model predicts that reduced cell growth on ethanol was a result of a disruption in metabolism leading to a reduced capacity to generate energy (Figure 4F). *In silico* rescue experiments identified 70 of the 248 perturbed enzymes as candidates that could rescue the ethanol-phase growth defect of SY14 (Supplemental Table II). Some of these genes (7/70) were deleted during chromosome fusion and were members of multicopy gene families whose individual contributions to metabolism are unclear. The remaining genes (63/70) were downregulated metabolic enzymes whose coding sequences were not perturbed in SY14, like *ACS1, PDH1*, and *YAT1*. The 70 rescue candidate genes were enriched amongst GO-slim terms related energy generation, including lipid metabolism, nucleotide metabolism, and carbohydrate metabolism (Figure 4G), consistent with our model of SY14 showing a growth defect using ethanol as the carbon source in Figure 4F.

## Discussion

In this work we investigated an *S. cerevisiae* strain whose genome is packaged into a single chromosome (strain SY14) and observed diminished growth on ethanol compared to a reference strain. Relatively few differentially expressed genes were observed in SY14 compared to reference during growth on ethanol, but the genes that were less expressed were enriched for GO terms associated with non-fermentable carbon source growth. This was surprising as the majority of these genes had unaltered coding sequences, promoters, terminators, as well as upstream and downstream genes in SY14 compared to reference. Metabolic models that simulated decreased flux through the enzymes that correspond to the under-induced genes (e.g. Ald3, Acs1, and Pdh1) predicted diminished ATP generation for SY14 on ethanol, which may explain diminished biomass accumulation. The exometabolite analysis herein suggests that a bottleneck to ethanol catabolism downstream of Alcohol Dehydrogenase, as ethanol depletion from media was similar in SY14 compared to the reference strain. These observations predict an accumulation of intermediates of ethanol catabolism in SY14, some of which (e.g. acetaldehyde) are toxic^26^ and could cause the observed ethanol sensitivity.

Previous characterization of SY14 found that the genome exhibited disrupted interchromosomal interactions and more globular chromosome structures compared to wildtype chromosomes^13,27^. Despite this disruption in chromosome interactions and topology, our analysis found that less than 2% of genes were differentially expressed in SY14 compared to reference. These observations suggest that interchromosomal interactions and topology are not strong influencers for the majority of genes in *S. cerevisiae* for the conditions tested. Instead, we hypothesize that these forces might influence specific genes, such as those activated as cultures switch from glucose to non-fermentable carbon sources^28^. This could explain the switch-like change in gene expression that coincides with observed changes in genome organization that occur post-diauxic shift^30^. SY14’s large chromosome may be too constrained to achieve this reorganization, which might influence a gene expression regime change. We propose that the sixteen chromosomes of *S. cerevisiae* enable greater plasticity than a single large chromosome, which may facilitate more dynamic chromosomal reorganization and gene regulation that is important during ethanol-phase growth.

The single chromosome^13^ and two chromosome^14^ yeast strains provide new opportunities to understand yeast physiology in relation to chromosome number and size. In this work, we found that the single chromosome yeast strain exhibits diminished ethanol tolerance and metabolism, which are key adaptations that enable *S. cerevisiae* to thrive and compete for sugar-rich niches as a Crabtree-positive yeast^28^. Notably, ethanol tolerance remains poorly understood and is a key limitation to industrial bioethanol fermentation, which utilizes *S. cerevisiae* strains to produce approximately 100 billion liters of ethanol annually^30^. Future efforts to repair diminished ethanol tolerance of SY14 via adaptive laboratory evolution and/or analysis of the other chromosomal fusion strains generated by Shao et al. (those with 2-15 chromosomes)^13^ may inform the design of improved strains for biotechnology. In addition, SY14 may be a valuable experimental system to understand chromosome number selection in yeast and may provide critical insights to inform the engineering of designer yeasts with synthetic genomes, as has been achieved in prokaryotes^31^ and viruses^32^.

## Methods

### Strains and Cultivation Conditions

The wildtype (BY4742) and single chromosome strain (SY14) were acquired from the lab of Zhongjun Qin and were grown at 30°C throughout this work. The batch fermentations in Figure 1 were carried out in YPD media with 1% glucose in a 500mL working volume bioreactor. Strains in Figure 2 and in Supplemental Figures were cultivated in YPD with 2% glucose (liquid media), or YP agar with 2% glucose, 3% glycerol, 3% ethanol or 6% ethanol. For Supplemental Figure 5, the CIT3 ORF was expressed from a klURA3 marked 2um plasmid flanked by the endogenous CIT3 promoter and terminator. A control plasmid was constructed from the aforementioned construct by removing the CIT3 ORF.

### Analysis of Doubling Time, Lag Phase, and Final OD

Doubling times were calculated using a non-linear fit of the exponential phase of glucose growth for each strain. This analysis was based on CO_2_ evolution for Figure 1 and OD measurements for Figure 2, and Supplemental 2. Lag-phase measurements were defined as the time elapsed for the first 1.5 doublings, which represents the time between the initial inoculation at 0.1OD_600_ to the cultures reaching 0.25OD_600_. Final OD was measured after five days of growth.

### Exometabolite measurements

Extracellular metabolites including glucose, ethanol, glycerol, pyruvate, and acetate, were quantified using an HPLC system (ultimate 3000 HPLC, Thermo Fisher) with a BioRad HPX-87H column (BioRad) and an IR detector, with 5 mM H_2_SO_4_ as the elution buffer at a flow rate of 0.6 mL/min and an oven temperature of 45 °C.

### Collection and Analysis of RNAseq Data

Biomass for RNAseq was collected in mid-glucose phase (7.5hours after inoculation) and during ethanol phase (20hours after inoculation). RNA extractions were performed on samples that were mechanically lysed with 0.5mm acid washed beads using an MP-Biomedicals™ FastPrep-24 for three one-minute cycles. Further extraction was performed using an RNeasy® Kit from Qiagen. Libraries were prepared using the TruSeq mRNA Stranded HT kit. Sequencing was carried out using an Illumina NextSeq 500 High Output Kit v2 (75 bases), with a minimum of 8 million paired-end reads per replicate. The Novo Nordisk Foundation Centre for Biosustainability (Technical University of Denmark), performed the RNA sequencing and library preparation. RNAseq was mapped with STAR and reads were assigned with featureCounts. Differential expression results were generated using scripts from the OrthOmics pipeline (https://github.com/SysBioChalmers/OrthOmics) from Doughty et al. 2020^33^, which is based on the limma and edgeR R packages. Raw datasets were uploaded to SRA under the accession number PRJNA594518 and differential expression results are reported in Supplemental Table I. Gene Ontology analysis was performed with the R-package Piano.

### Metabolic Modeling

The genome-scale metabolic model ecYeast 9.3^24^ was used to generate enzyme constrained models for both reference (BY4742) and SY14 strains using the GECKO toolbox^25^. The default enzyme pool parameter of 0.1 was used as the upper limit for enzyme abundance. To model glucose growth, glucose was set as the sole carbon source, and exchange fluxes such as glucose uptake rate, by-product production rates, and biomass formation rate were constrained to observed values for both reference and SY14 strains. The non-growth associated maintenance energy (NGAM) reaction for each model was set as the objective function, and flux balance analysis (FBA) was used to calculate the maximum NGAM for both reference and SY14 strains. To model ethanol growth, we constructed models for both reference and SY14 strains as follows: first, the default ecYeast9.3 model was constrained with ethanol as the sole carbon source and growth rate constrained to 0.22 h^-1^, with a flexibilization factor of ±5%. We then performed random sampling with a pair of randomly weighted objective functions to obtain a set of 1,000 feasible flux distributions in reference strain cell growth^34^. Then, for each flux distribution for the reference strain, we constrained the upper bound of enzyme exchange reactions of the 248 differentially expressed enzymes in SY14 by multiplying the simulated enzyme usage in the reference strain model with the fold-change value in gene expression analysis, with a flexibilization factor of ±20%. The objective function was set first to maximize growth rate; infeasible solutions (4 out of 1,000) were discarded. Then, with the growth rate constrained to the maximum calculated value with a flexibilization factor of ±5%, the objective function was set to maximize both ethanol consumption and NGAM. For the *in silico* rescue experiment, we removed the constraints on the 248 enzymes one at a time for each of the 996 feasible solutions and used FBA to calculate the maximum growth rate using ethanol as the sole carbon source. The average maximum growth rate was calculated, and the 70 enzymes that rescued the mean growth rate to 0.22 h^-1^ with a flexibilization factor of ±5% are subjected to GO-slim enrichment analysis at Saccharomyces genome database (https://www.yeastgenome.org/). Enriched GO-slim terms with >5 genes were included.

## Contributions

T.W.D. collected and analyzed RNAseq and genomics, performed batch fermentations, and wrote the manuscript. R.Y. performed HPLC, qPCR, cell morphology assays, and constructed metabolic models. L.F.C. performed growth assays and designed/built/tested CIT3 plasmids. Z.Q., V.S., and J.N. conceived and supervised the project.

## Acknowledgements

This project has received funding from the European Union’s Horizon 2020 Framework Programme for Research and Innovation -Grant Agreement No. 720824. This work was also supported by the Knut and Alice Wallenberg Foundation and the Novo Nordisk Foundation (Grant no NNF10CC1016517).

## References

1. Stinchcomb, D. T., Struhl, K. & Davis, R. W. Isolation and characterisation of a yeast chromosomal replicator. Nature 282, 39–43 (1979).

2. Clarke, L. & Carbon, J. Isolation of a yeast centromere and construction of functional small circular chromosomes. Nature 287, 504–509 (1980).

3. Blackburn, E. H. Artificial chromosomes in yeast. Trends Genet. 1, 8–12 (1985).

4. Koshland, D. E., Mitchison, T. J. & Kirschner, M. W. Polewards chromosome movement driven by microtubule depolymerization in vitro. Nature 331, 499–504 (1988).

5. de Lange, T. How Telomeres Solve the End-Protection Problem. Science (80-.). 326, 948 LP – 952 (2009).

6. Sanger, F. & Coulson, A. R. A rapid method for determining sequences in DNA by primed synthesis with DNA polymerase. J. Mol. Biol. 94, 441–448 (1975).

7. Murray, A. W. & Szostak, J. W. Construction of artificial chromosomes in yeast. Nature 305, 189–193 (1983).

8. Szostak, J. W. & Blackburn, E. H. Cloning yeast telomeres on linear plasmid vectors. Cell 29, 245–255 (1982).

9. Nagalakshmi, U. et al. The Transcriptional Landscape of the Yeast Genome Defined by RNA Sequencing. Science (80-.). 320, 1344 LP – 1349 (2008).

10. Lieberman-Aiden, E. et al. Comprehensive Mapping of Long-Range Interactions Reveals Folding Principles of the Human Genome. Science (80-.). 326, 289 LP – 293 (2009).

11. Shendure, J. & Ji, H. Next-generation DNA sequencing. Nat. Biotechnol. 26, 1135–1145 (2008).

12. Cong, L. et al. Multiplex Genome Engineering Using CRISPR/Cas Systems. Science (80-.). 339, 819 LP – 823 (2013).

13. Shao, Y. et al. Creating a Functional Single-Chromosome Yeast. Nature 560, 331–335 (2018).

14. Luo, J., Sun, X., Cormack, B. P. & Boeke, J. D. Karyotype engineering by chromosome fusion leads to reproductive isolation in yeast. Nature 560, 392–396 (2018).

15. Gordon, J. L., Byrne, K. P. & Wolfe, K. H. Mechanisms of chromosome number evolution in yeast. PLoS Genet. 7, e1002190–e1002190 (2011).

16. Neurohr, G. et al. A Midzone-Based Ruler Adjusts Chromosome Compaction to Anaphase Spindle Length. Science (80-.). 332, 465 LP – 468 (2011).

17. Cremer, T. & Cremer, C. Chromosome territories, nuclear architecture and gene regulation in mammalian cells. Nat. Rev. Genet. 2, 292–301 (2001).

18. Ji, X. et al. 3D Chromosome Regulatory Landscape of Human Pluripotent Cells. Cell Stem Cell 18, 262–275 (2016).

19. Jiang, S. & Dai, J. Inevitability or contingency: how many chromosomes do we really need? Sci. China Life Sci. 62, 140–143 (2019).

20. Taddei, A. et al. The functional importance of telomere clustering: Global changes in gene expression result from SIR factor dispersion. Genome Res. 19, 611–625 (2009).

21. Graybill, E. R., Rouhier, M. F., Kirby, C. E. & Hawes, J. W. Functional comparison of citrate synthase isoforms from S. cerevisiae. Arch. Biochem. Biophys. 465, 26–37 (2007).

22. Giaever, G. et al. Functional profiling of the Saccharomyces cerevisiae genome. Nature 418, 387–391 (2002).

23. Monteiro, P. T. et al. YEASTRACT+: a portal for cross-species comparative genomics of transcription regulation in yeasts. Nucleic Acids Res. 48, D642–D649 (2019).

24. Lu, H. et al. A consensus S. cerevisiae metabolic model Yeast8 and its ecosystem for comprehensively probing cellular metabolism. Nat. Commun. 10, 3586 (2019).

25. Sánchez, B. J. et al. Improving the phenotype predictions of a yeast genome-scale metabolic model by incorporating enzymatic constraints. Mol. Syst. Biol. 13, 935 (2017).

26. Aranda, A. & del Olmo, M. Exposure of Saccharomyces cerevisiae to Acetaldehyde Induces Sulfur Amino Acid Metabolism and Polyamine Transporter Genes, Which Depend on Met4p and Haa1p Transcription Factors, Respectively. Appl. Environ. Microbiol. 70, 1913 LP – 1922 (2004).

27. Duan, Z. et al. A three-dimensional model of the yeast genome. Nature 465, 363–367 (2010).

28. Dashko, S., Zhou, N., Compagno, C. & Piškur, J. Why, when, and how did yeast evolve alcoholic fermentation? FEMS Yeast Res. 14, 826–832 (2014).

29. Guidi, M. et al. Spatial reorganization of telomeres in long-lived quiescent cells. Genome Biol. 16, 206 (2015).

30. Mohd Azhar, S. H. et al. Yeasts in sustainable bioethanol production: A review. Biochem Biophys Rep 10, 52–61 (2017).

31. Gibson, D. G. et al. Complete Chemical Synthesis, Assembly, and Cloning of a Mycoplasma genitalium Genome. Science (80-.). 319, 1215 LP – 1220 (2008).

32. Blight, K. J., Kolykhalov, A. A. & Rice, C. M. Efficient Initiation of HCV RNA Replication in Cell Culture. Science (80-.). 290, 1972 LP – 1974 (2000).

33. Doughty, T. W. et al. Stress-induced expression is enriched for evolutionarily young genes in diverse budding yeasts. Nat. Commun. 11, 2144 (2020).

34. Bordel, S., Agren, R. & Nielsen, J. Sampling the solution space in genome-scale metabolic networks reveals transcriptional regulation in key enzymes. PLoS Comput. Biol. 6, e1000859–e1000859 (2010).

