## Supplemental Figures for "A Single Chromosome Strain of *S. cerevisiae* Exhibits Diminished Ethanol Metabolism and Tolerance"

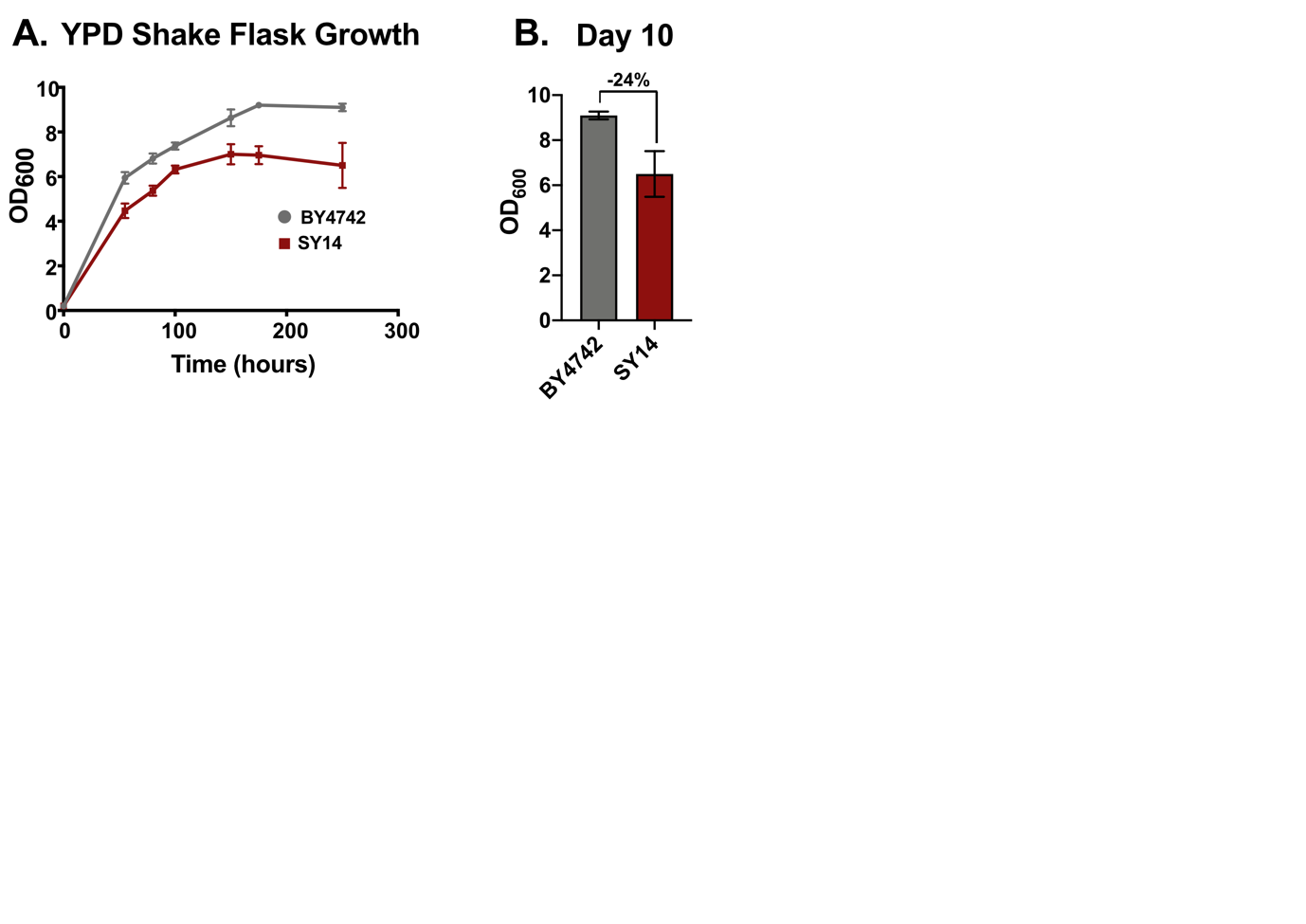

**Supplemental Figure 1: Long-term shake flask growth of SY14 results in diminished biomass accumulation.** BY4742 and SY14 were grown in YPD media in shake flasks for 10 days. Measurements were taken periodically and are shown in A. The difference in biomass on day 10 was 24% (B).

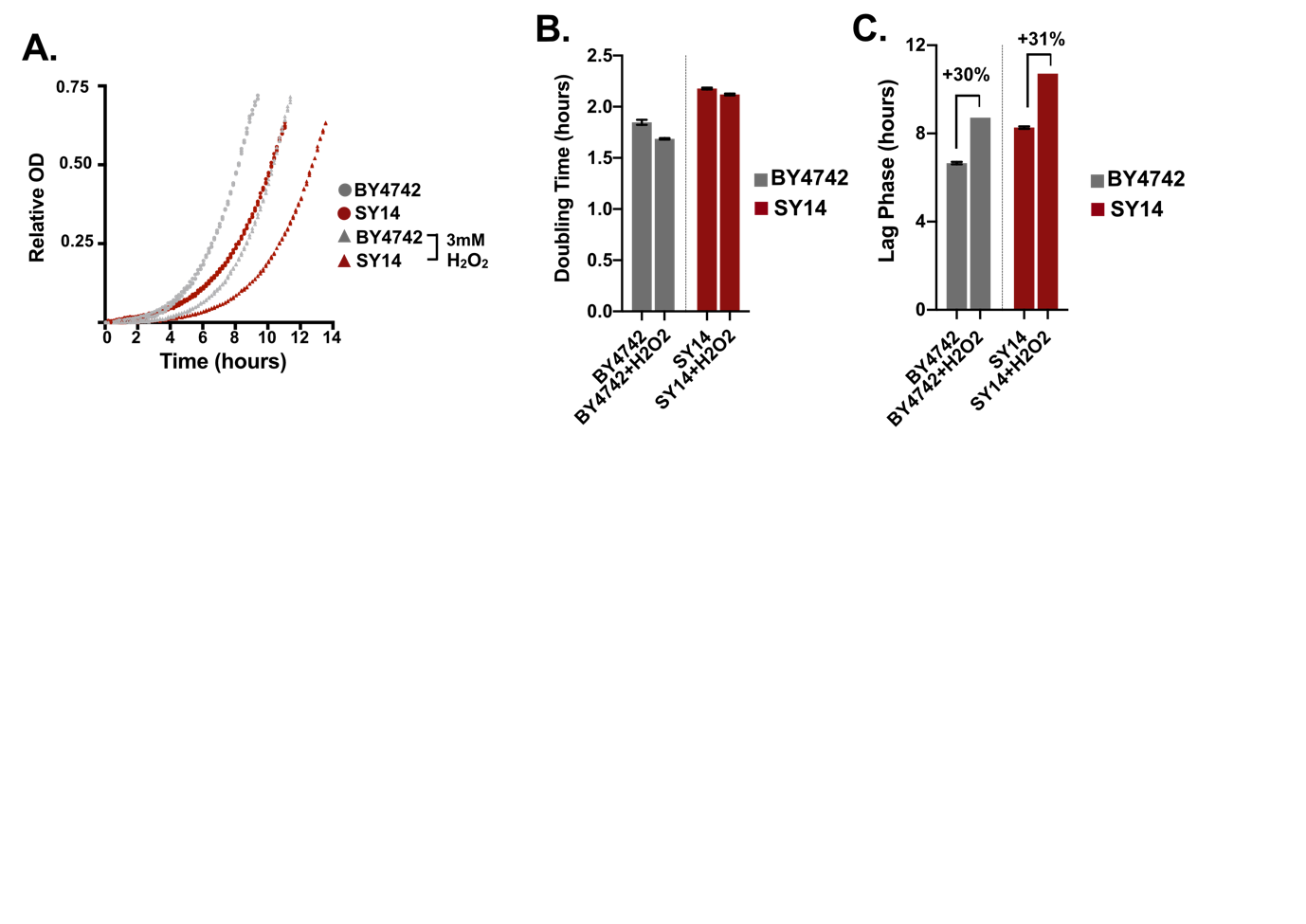

**Supplemental Figure 2: SY14 does not exhibit increased sensitivity to Hydrogen Peroxide.** A. Growth curves were calculated using a 48-well growth assay measuring OD_600_ every 10minutes for wildtype (BY4742) and single chromosome (SY14) strains. The maximum doubling time during glucose phase (B) and lag phase (C) are shown.

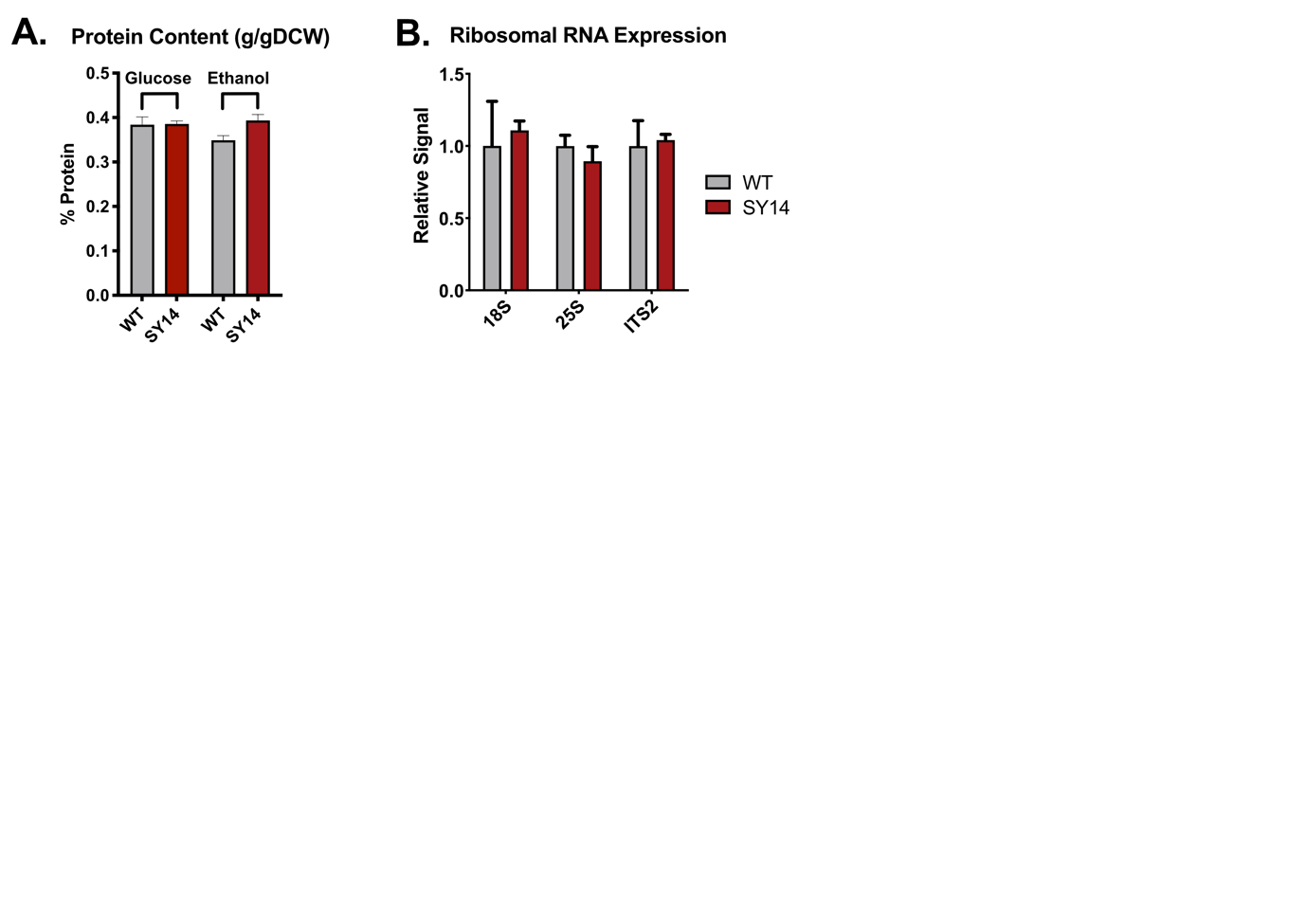

**Supplemental Figure 3: Total protein and rRNA abundance and processing are similar between wildtype and SY14 strains.** A. Protein content was measured via Lowry assay. B. To assess potential changes in Ribosomal RNA expression or processing, rRNA was measured via qPCR for mature rRNA regions (18S and 25S), as well as a region that is removed during maturation (ITS2).

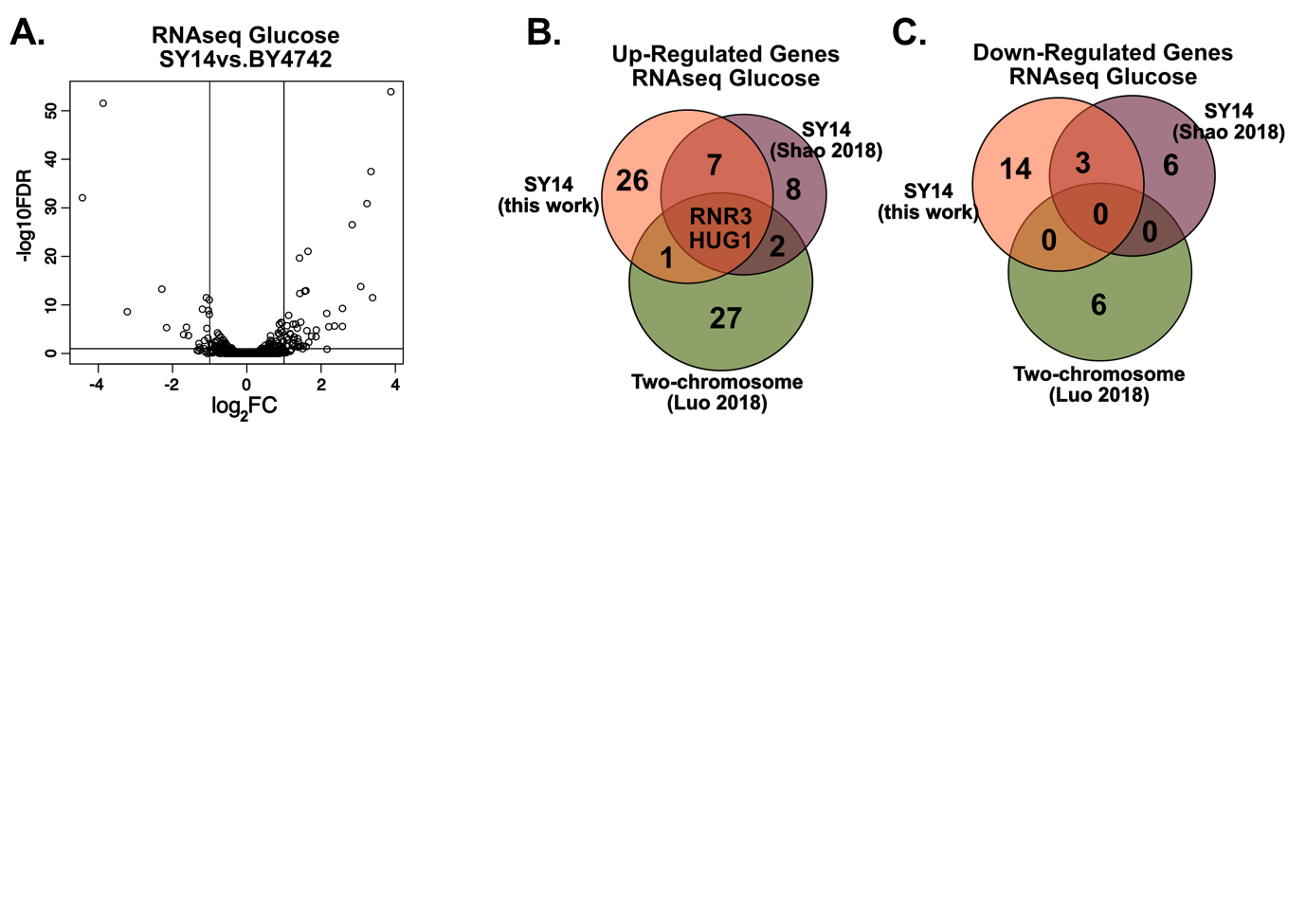

**Supplemental Figure 4: Glucose RNAseq suggests RNR3 and HUG1 are differentially expressed in chromosomal fusion strains grown on glucose.** A. Glucose-phase differential expression data was generated using biomass from fermentations shown in Figure 1. B. The data in this report (red) was compared to RNAseq from previous reports that studied either the single chromosome strain (purple) or the two chromosome strain (green). Shared differentially expressed genes (log_2_FC>1_abs_ FDR<0.01) were compared amongst up (B) and downregulated (C) genes.

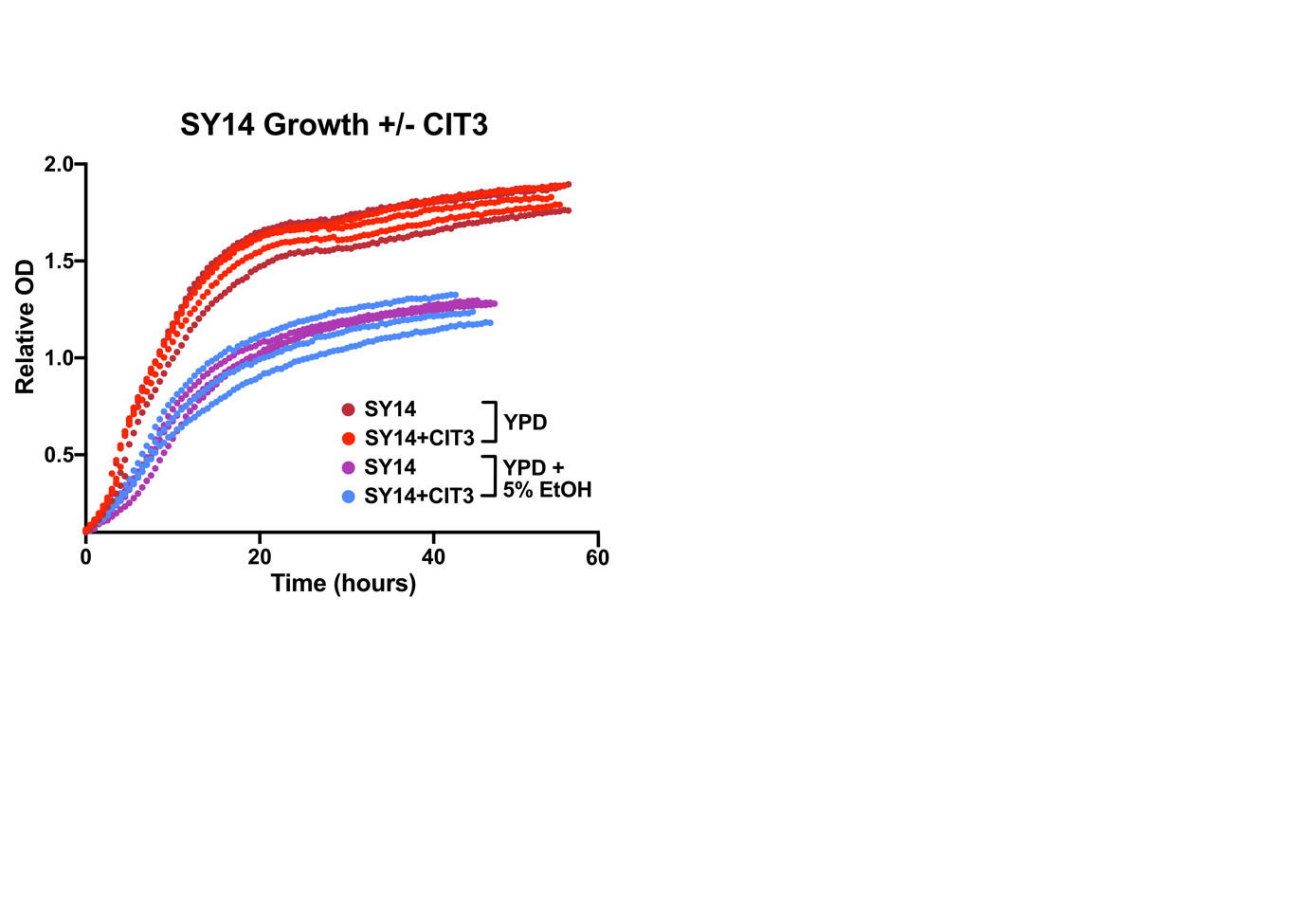

**Supplemental Figure 5: A plasmid borne copy of CIT3 does not rescue SY14 growth rate or ethanol sensitivity.** A plasmid with the CIT3 promoter and terminator (brown and purple) or plasmid with CIT3 promoter, ORF, and terminator (dark red and light blue) were transformed into SY14. Growth was monitored in 48-well plate format by measuring OD_600_ every 10 minutes in the presence of absence of ethanol.

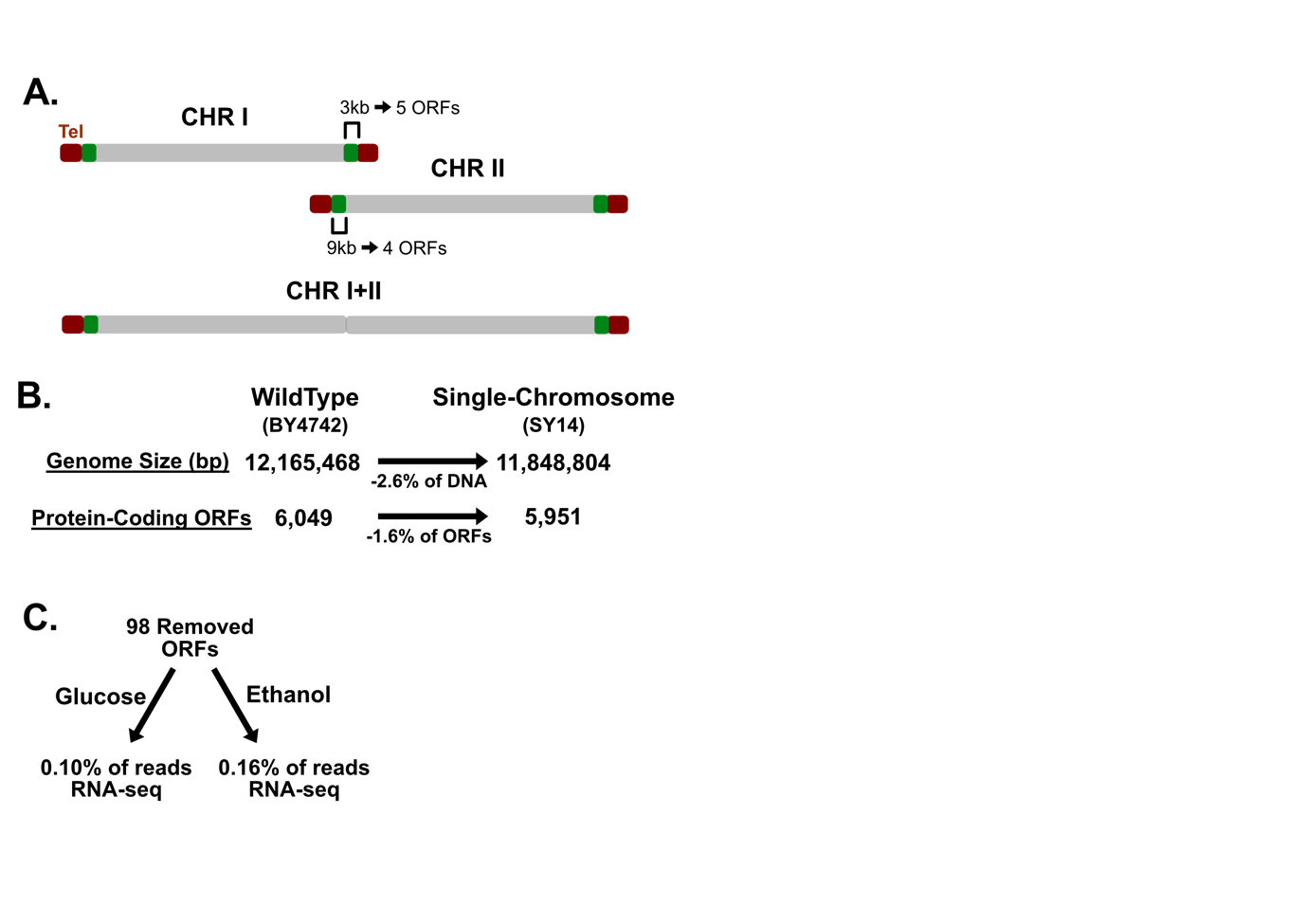

**Supplemental Figure 6: SY14 Deleted Genes are evolutionarily young genes that are poorly expressed.** A. Chromosome I and II fusion is shown as an example of the cause of ORF removal during the construction of SY14. B. Genome size and ORF count for wildtype (BY4742) and single chromosome (SY14) strains. C. The removed ORFs in SY14 were queried for the % of total ORF-associated reads for BY4742 during glucose and ethanol growth.

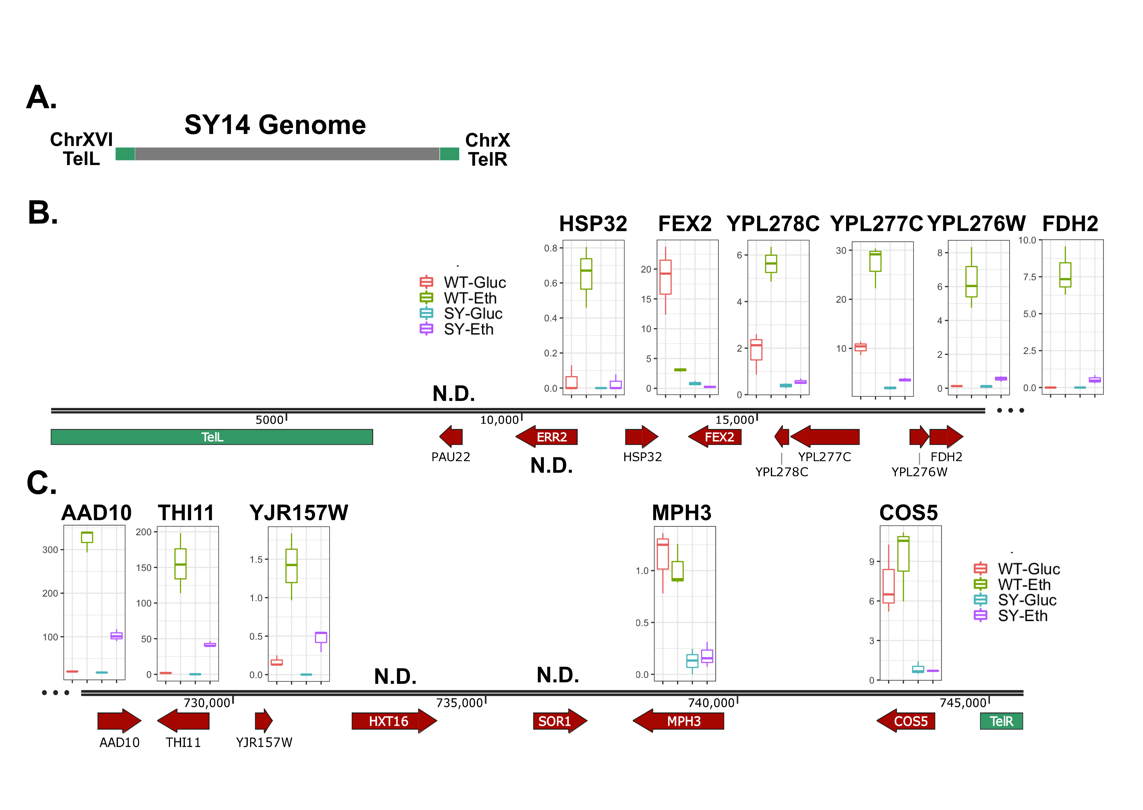

**Supplemental Figure 7: Subtelomeric genes are misregulated in SY14.** A. The single chromosome in strain SY14 is shown. Genes detected in our RNAseq analyses are shown for the left telomere (B) and the right telomere (C).
